# Assessment of Contralateral Efferent Effects in Human *Via* ECochG

**DOI:** 10.1101/2024.12.24.630246

**Authors:** Eric Verschooten, Elizabeth A. Strickland, Nicolas Verhaert, Philip X. Joris

## Abstract

Efferent projections from the brainstem to the inner ear are well-described anatomically and physiologically but their precise function remains debated. The medial olivocochlear (MOC) system and its reflex, the MOCR, have been particularly well studied. In animals, anatomical and physiological data are fine-grained and extensive and suggest an important role for the MOCR in anti-masking e.g. to improve the detection of tones in background noise. Extensive behavioral studies in human support this role, but direct linking of behavioral paradigms to the MOCR is challenging because of the difficulty in obtaining appropriate human neural measures. We developed a new approach in which mass potentials were recorded near the cochlea of normal hearing and awake human volunteers to increase the signal-to-noise (SNR) ratio, and examined whether broadband noise to the contralateral ear elicited MOCR anti-masking effects as reported in animals. Probing the mass potential to the onset of brief tones at 4 and 6 kHz, convincing anti-masking or suppressive effects consistent with the MOCR were not detected. We then changed the recording technique to examine the neural phase-locked contribution to the mass potential in response to long, low-frequency tones, and found that contralateral sound suppressed neural responses in a systematic and progressive manner. We followed up with psychophysical experiments in which we found that contralateral noise elevated detection threshold for tones up to 4 kHz. Our study provides a new way to study efferent effects in the human peripheral auditory system and shows that contralateral efferent effects are biased towards low frequencies.

## Introduction

One of the intriguing organizational properties in hearing and balance is that the end organs are under control by the brain. The anatomical organization of efferent projections to the inner ear is relatively straightforward and well-described, but it has been surprisingly difficult to formulate a precise *raison d’être* for these projections (Lauer et al., 2022; Mackrous et al., 2022). Most studies of efferent effects in hearing (Robles and Delano, 2008; Guinan, 2011; Fuchs and Lauer, 2019; Jennings, 2021) have focused on the anti-masking properties of the medial olivocochlear (MOC) system. Simply put, the detection of a sound can be hampered by the presence of other sounds, and activation of the MOC reflex (MOCR) can reduce such masking. MOC efferents inhibit their target, the outer hair cells. In ill-understood ways, outer hair cells increase cochlear gain. Thus, activation of the MOCR reduces cochlear gain. Somewhat counterintuitively, by reducing cochlear sensitivity, MOCR activation can reduce masking. A functional parallel is found in the middle ear muscle reflex (MEMR), which also reduces cochlear activation albeit through entirely different (conductive) mechanisms. MEMR and MOCR operate over different frequency ranges: the MEMR reduces the effectiveness of low-frequency maskers and the MOCR counteracts masking of transients in high-frequency signals (Liberman and Guinan, 1998).

Electrophysiological studies in experimental animals have enabled characterization of the MOC system in great detail. To study humans, investigators have mainly used two non-invasive techniques. Psychophysical stimulus paradigms and procedures have been designed to specifically reveal efferent effects. Their attraction is the potential to address the role of efferents in human perception, but the causal role of efferents in the effects measured is necessarily indirect. Studies of otoacoustic emissions (OAEs) in humans complement behavioral measurements by providing a direct and noninvasive window on the cochlea, but suffer from technical constraints (e.g. in the frequency range of measurements) and their unclear relationship to neural cochlear output (Puria et al., 1996). An extensive review of the role of efferents in human hearing concluded that “innovative approaches are needed to resolve the dissatisfying conclusion that current results are unable to definitely confirm or refute the role of the MOC reflex” (Jennings, 2021).

Only limited attempts have been made to apply the electrophysiological methods used in animals to humans. Recordings from single neurons are obviously out-of-reach, but a vast corpus of efferent work in experimental animals has relied on cochlear mass potentials recorded at the middle ear. In animals, within-subject comparisons shows stronger efferent effects in neural mass potentials than in OAEs (Puria et al., 1996). Electrodes in the human outer ear, e.g. on the tympanic membrane, have been used toward electrophysiological study of efferent effects (for references: see Discussion), but the tiny signal-to-noise ratio (SNR) in such recordings severely limits their power. Previous work in our laboratory has made use of a minimally invasive procedure using a transtympanic approach (Verschooten and Joris, 2022) to record cochlear potentials from normal-hearing volunteers, enabling us to address issues of frequency tuning and temporal coding (Verschooten et al., 2018). This technique provides an avenue to apply stimulus paradigms previously used in animal and psychophysical studies of the MOCR and thereby the potential to bridge the two.

The MOC system is bilateral so that sound in one ear can trigger efferent effects in the same (ipsilateral) ear and in the other (contralateral) ear. In a previous study using transtympanic recording (Verschooten et al., 2017), we reported on effects of the ipsilateral MOCR. Here we study the contralateral MOCR, using both transtympanic physiological recordings and psychophysical measurements.

## Results

Early electrophysiological work in human showed a large effect of contralateral stimulation on the neural response to a 4 kHz probe tone (Folsom and Owsley, 1987), which is also the probe frequency most commonly studied in human behavioral studies, because large effects were found at this frequency in studies of temporal effects in simultaneous masking (Krull and Strickland, 2008). We first physiologically assessed the contralaterally-elicited MOCR with tone pips at 4 and 6 kHz in 5 subjects and found that the effects were small and variable. We widened our search for contralaterally-elicited MOCR effects using a non-masking paradigm and a different assessment method (neurophonic) and found consistent suppressive effects at low frequencies (≤ 800 Hz). We then assessed the effects of contralateral stimulation using a behavioral paradigm and found effects over a frequency range < 4 kHz.

### Electrophysiological suppressive and anti-masking effects at 4-6 kHz

We illustrate anti-masking effects in 2 extensively tested subjects (Figs. 2-4). To assess the anti-masking effect of the MOCR, a large number of stimulus parameters need to be chosen for the ipsilateral probe tone and masker as well as the contralateral elicitor. The stimulus parameters we selected were guided by previous physiological studies in animals and psychophysical studies in human, but were constrained by the SNR of the recordings. To determine an appropriate level of the probe tone, we first ran intensity series using paradigm SP0 (Fig. 1).

**Figure 1.**
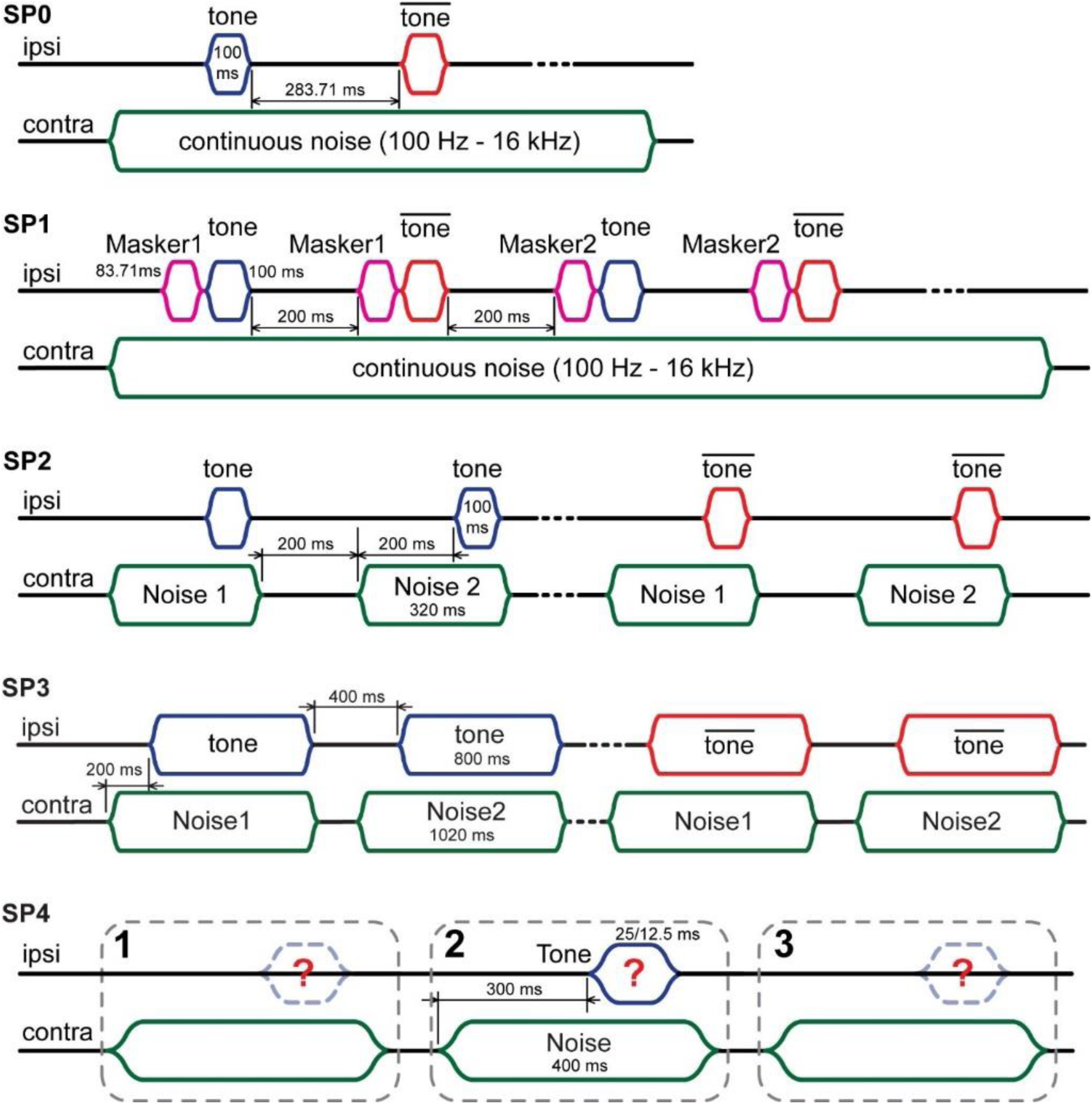
Timelines illustrating the different stimulus paradigms. SP0 to SP3 were used in electrophysiological experiments. SP0 and SP1 were designed to investigate the contralateral anti-masking effect; SP2 and SP3 to assess the contralateral influence on the CAP (SP2), and on the CM and neurophonic at low frequencies (SP3). SP4 was the paradigm for the 3AFC method used in the psychoacoustical experiments. Red and blue envelopes indicate the pure-tone probe tone at the experimental (ipsilateral) ear. Green envelopes indicate the broadband noise elicitor at the contralateral ear. Magenta envelopes indicate the ipsilateral forward masker.

**Figure 2.**
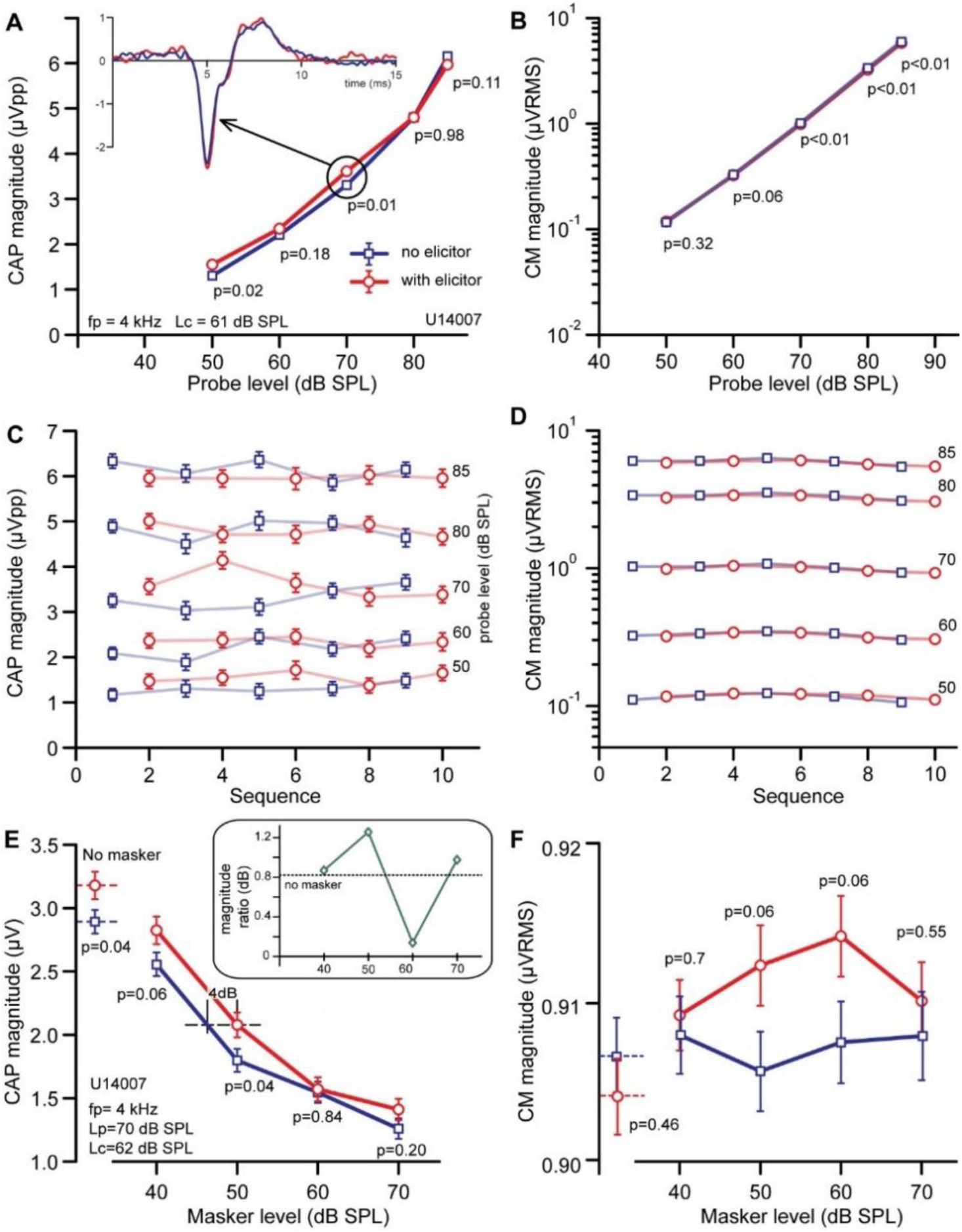
Unmasking effects in one subject. A-D: data without masking, to determine suitable level of the 4 kHz probe. A: CAP response to unmasked probe presented at different sound levels with (red) and without (blue) contralateral elicitor. The elicitor was a continuous contralateral noise at 61 dB SPL (= Lc). Inset shows CAPs at 70 dB, using the same color code. B: CM extracted from same responses. C, D: Sequential CAP (C) and CM (D) amplitudes for the different runs in chronological order, underlying average values of A and B. Lines connect responses with same probe level. Odd runs are without elicitor, even with elicitor. E, F: Effect of contralateral elicitor on CAP (E) and CM (F) to a probe at 4 kHz in the presence of a forward masker at a range of sound levels. The data closest to the ordinate are for the no-masker condition. The inset in E shows the ratio (dB) for the data with and without elicitor; the dashed line is the ratio of the no-masker condition. P-values are calculated under H0: μ1 = μ2. fp = probe frequency, Lp = probe level, Lc = elicitor level. Notice compressed y-scale in F. Error bars indicate SEM: obtained in C, D by bootstrapping and in A, B, E, and F by propagated SEM.

A 4-kHz probe-tone was varied in intensity (50 – 85 dB) in the presence or absence of a sustained contralateral elicitor. CAP (Fig. 2A,C) and CM (Fig. 2B,D) measurements are shown for one subject, with (red circles) and without (blue squares) contralateral elicitor at 61 dB SPL. Both the CAP (Fig 2A) and CM (Fig. 2B) increase monotonically with probe level. At the lower SPLs, the CAP amplitude is somewhat larger when an elicitor is present (i.e. an enhancement), with a significant difference at 50 and 70 dB SPL (Fig. 2A). The corresponding CM, plotted on a log-log scale (Fig. 2B), is linearly dependent on probe level, and shows a very small decrease in the presence of the elicitor, significant for the three highest levels (70-85 dB SPL). To check whether there is a systematic pattern over time, Fig. 2C, D illustrates CAP and CM magnitudes for the sequence of measurements, where each measurement block (each symbol) represents somewhat less than 7 minutes of recording. The mean CAP values and variability are reasonably stable and the difference between elicitor and no-elicitor conditions is quite consistent over the first couple of sessions but appears to decrease over time (Fig. 2C). The CM has small variance and is stable across measurement blocks but shows a small and slow up-then downward drift (Fig. 2D).

Opposite effects of efferent activation on CAP and CM are seen as a hallmark of efferent activation (e.g., (Elgueda et al., 2011)), but the increase in CAP and decrease in CM observed in Fig. 2A-D are opposite to the expected pattern. Nevertheless, the data are not necessarily inconsistent with an efferent effect. The “classical” increase in CM is variable and not always observed (Guinan, 2006; Aedo et al., 2015), and the decrease observed here is very small. Also, an in-rather than a decrease in CAP in response to a contralateral elicitor is sometimes observed in animal studies (Kawase and Liberman [1993], see Discussion). Based on the data in Fig. 2A-D, a probe tone level of 70 dB SPL was chosen to test the anti-masking effect of a contralateral elicitor. The SNR at this level is sufficient given the limited measurement time and the level is adequate to expect an anti-masking effect, based on data in cat.

Paradigm SP1 (Fig. 1) was used to assess the anti-masking effect of the MOCR. This is similar to paradigm SP0 except that now the ipsilateral pure-tone probe is preceded by a Gaussian noise masker, which is varied in level. Fig. 2E, F shows magnitude of CAP and CM, respectively, for different masker levels (none, 40, 50, 60, 70 dB SPL), with (red) and without (blue) contralateral elicitor. Data without the masker are shown on the far left near the ordinate. The elicitor causes a small increase in CAP (Fig. 2E), consistent with the effect already shown Fig. 2A for the same probe level. When a masker is added, the CAP decreases in amplitude with masker level (blue squares), as expected. Adding the elicitor (red circles) largely increases response amplitudes, i.e. a change in the same direction as for the unmasked response.

The changes are only significant at the lowest masker levels (40 and 50 dB SPL), and appear consistent with an MOCR anti-masking effect.

Forward masking hinges on temporal adaptation in synaptic transmission: it is first observed at the level of the auditory nerve and is not observed in receptor potentials. Therefore, the forward masker is not expected to affect the magnitude of the CM. Indeed, in the absence of the elicitor the CM is unaffected by the presence or level of the masker (Fig. 2F). The presence of the contralateral elicitor in the absence of a masker gave a small but non-significant suppression of the CM (consistent with Fig. 2D). Increase of the masker level caused a small but non-significant CM increase for masker levels of 50 and 60 dB SPL.

Overall, the results suggest a small anti-masking effect at low masker levels. A direct comparison between masker and no-masker conditions is provided by the inset in Fig. 2E, which shows the effect of the elicitor as a ratio, i.e. magnitude with elicitor (red) re. magnitude without elicitor (blue), expressed in dB. The condition without masker is provided as the dotted reference line at about 0.8 dB. Taking the data at face value, the anti-masking effect on CAP amplitude is ∼1 dB for all masker levels (including no masker) except at 60 dB. Expressed using an iso-output viewpoint (“effective attenuation”, see Puria et al. [1996] and Fig. 2E, horizontal dashed line), the increase in masker level required to keep the CAP magnitude constant when the elicitor is present (red) compared to when it is not (blue), is ∼4 dB. Effects of the masker on the CM are not significant and indicate an absence of the MEMR.

Fig. 3 shows similar measurements for another subject. While the effect on the CM (Fig. 3B) is qualitatively similar to that of the previous subject (Fig. 2B), here the CAP data (Fig. 3A) show a very small and non-significant reduction with a contralateral elicitor, in contrast to the increase seen in Fig. 2A. For the probe level at 70 dB SPL, the datapoints are nearly identical. A masking series (Fig. 3C) was run with the same conditions (probe frequency 4 kHz, level 70 dB) as for the previous subject, but at a 12-dB larger elicitor level. In this subject the elicitor causes a small reduction of the response to the unmasked probe (Fig. 3C, leftmost symbols) but has no significant effect when a masker is present. The CM responses (Fig. 3D) were an order of magnitude smaller than in the previous subject: the elicitor causes a reduction in CM amplitude but there is little effect of the masker.

**Figure 3.**
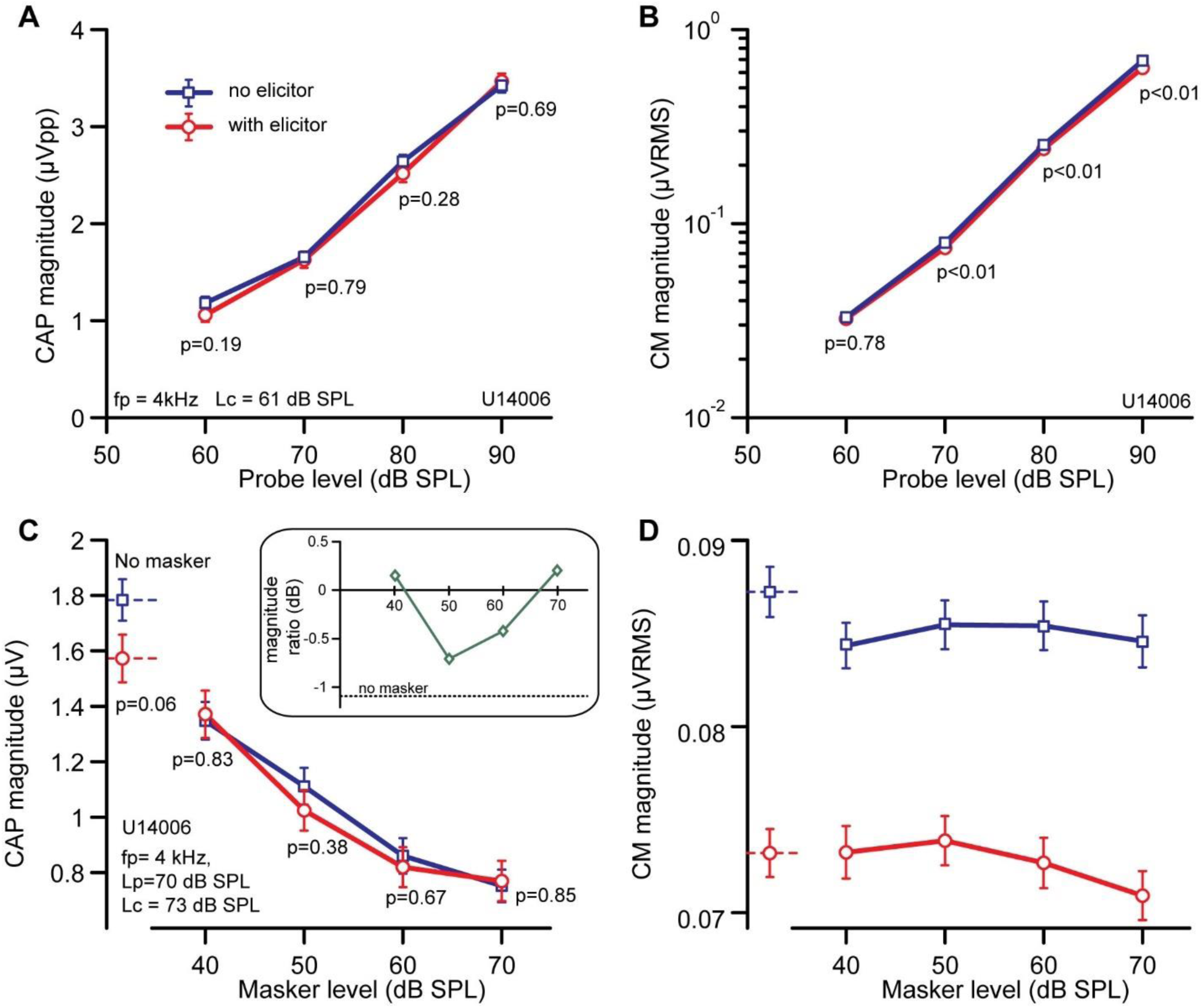
Effect of a continuous contralateral noise elicitor on a masked 4 kHz probe in a different subject. A,B,C,D are formatted as in Fig. 2A,B,E,F.

A full series of measurements was run again for this subject with a probe at 6 kHz (Fig. 4), for a wider range of probe and masker levels and in finer (5-dB) steps. The overall pattern of results for the probe level measurements (Fig. 4A,B) and masking series (Fig. 4C,D) is similar to that for the 4 kHz probe (Fig. 3). Across probe levels, the effect of the contralateral elicitor on CAP and CM is not significant (Fig. 4A,B). The same applies to the masking series data of CAP (Panel C) and CM (Panel D), for most data points. As with the other subject (Fig. 2E), the CAP values are a bit larger in the presence of elicitor and the absence of a masker, and this is also the case at the extreme levels of the masking function (i.e., 30, 70-80 dB SPL), resulting in positive ratios in the inset of Fig. 4C, but none of these changes is significant. The effects in CM (Fig. 4D) are consistent with Fig. 3D for the same subject and opposite to the first subject (Fig. 2F) and are again not significant.

**Figure 4.**
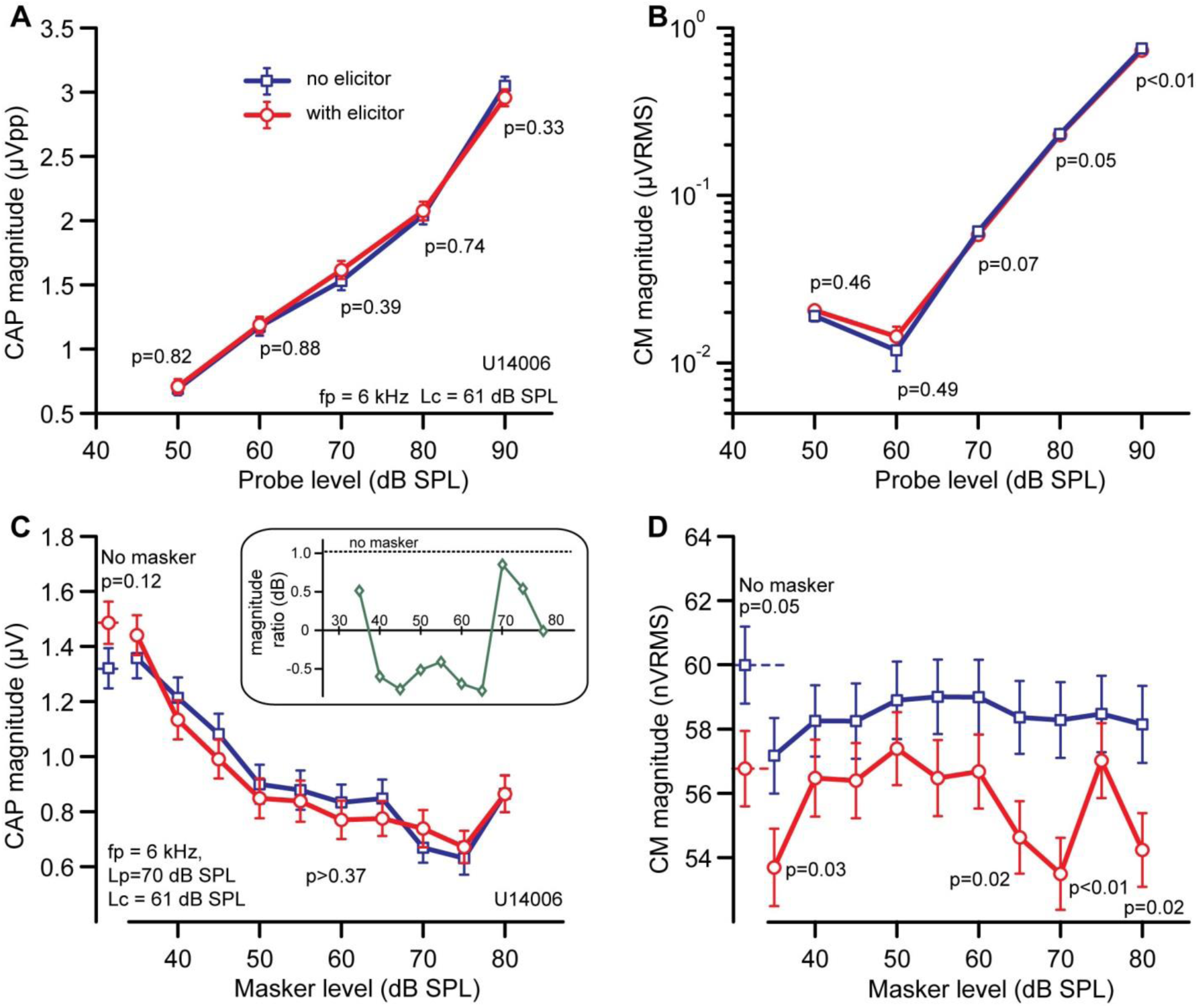
Contralateral effects for a probe tone of 6 kHz (same subjects as in Fig. 3). p-values (hypothesis H0: μ1 = μ2) in C were all >0.1, p-values in D are only stated when ≤0.1.

Given the experimental limitations of these human recordings in the face of the small and complex effects, inconsistent across subjects, we changed our approach in the subsequent experiments: rather than look for an anti-masking effect, we focused on a simpler paradigm that allowed us to explore a wider range of stimulus parameters and response metrics to zoom in on effects of a contralateral elicitor on responses to ipsilateral tonal probes.

### Electrophysiological effects of elicitor level and duration

The measurements described so far to study anti-masking effects made use of continuous presentation of the contralateral elicitor. We choose a continuous, broadband elicitor because animal experiments have shown that MOC efferents have minimal adaptation over both short and long timescales (Brown, 2001) and that a sustained broadband contralateral elicitor is more effective in inhibiting the response of auditory nerve fibers than a gated one (Warren and Liberman, 1989a). On the other hand, possibly the ipsilateral probe and masker trigger or sensitize efferents (Liberman, 1988; Guinan et al., 2003) and combine with slow contralateral effects (Larsen and Liberman, 2009) to result in sustained recruitment of the MOCR, which would also minimize differences in responses for conditions with and without elicitor. In any case, the lack of clear and consistent MOCR effects in the responses for paradigm SP1 invites further exploration of the ipsi- and contralateral parameters.

Using SP2 (Fig. 1), we presented an intermittent elicitor which could be varied in level. When time allowed, we also tested different levels of the probe tone. Fig. 5A shows CAP responses of one subject for 3 different probe levels (different colors) and a range of contralateral elicitors. Without contralateral elicitor (bold lines), the CAP amplitude increased with probe level from 50 to 60 dB SPL. There was little additional amplitude change from 60 to 70 dB SPL, but the latency of the response decreased with each increase in probe level. Responses to five different elicitor conditions (different line styles) are superimposed at each probe level. The elicitor caused small increases in amplitude of the CAP, most clearly visible in the prominent troughs, but note that these are not accompanied by clear latency changes.

**Figure 5.**
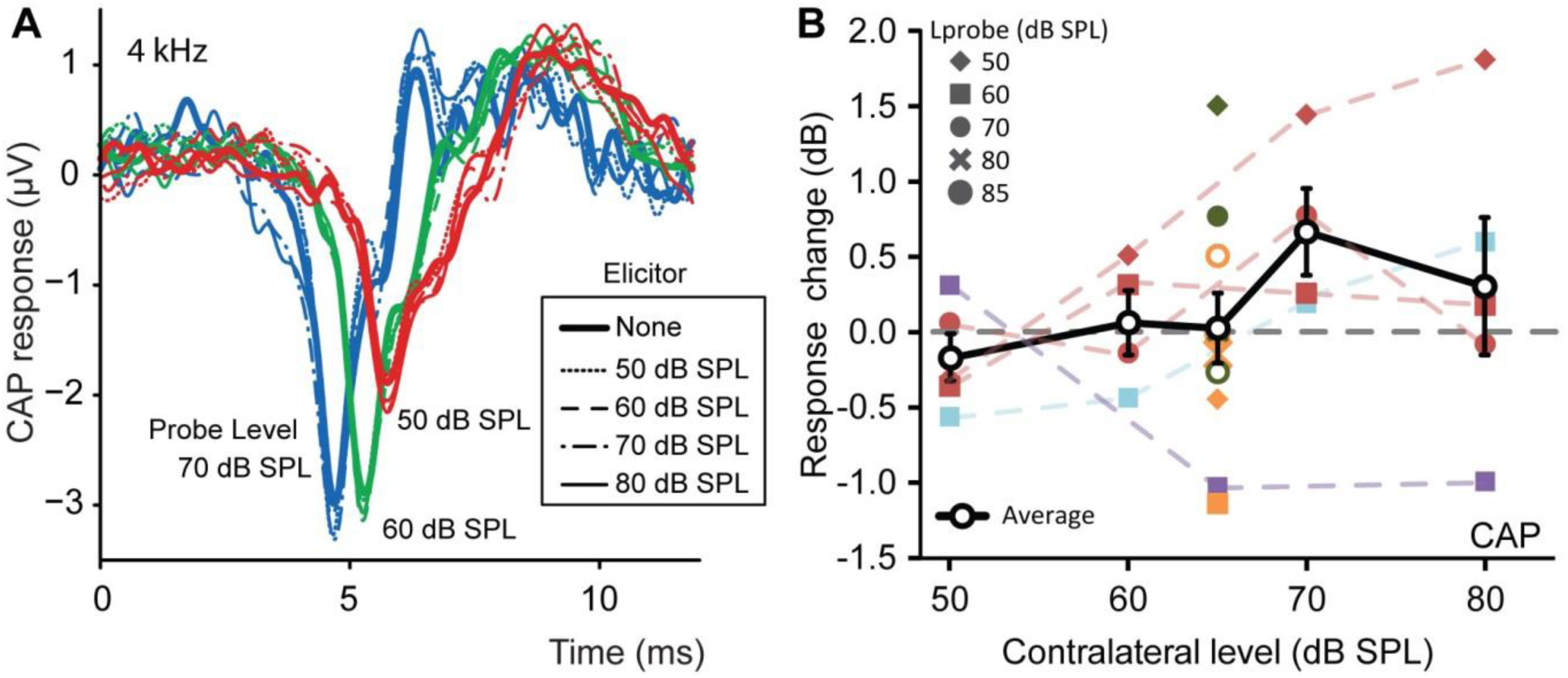
Influence of contralateral elicitors on an ipsilateral 4-kHz unmasked probe tone. A: Examples of CAP responses as a function of probe and elicitor level for one subject. Signals were denoised with a 0.05-4 kHz phase preserving band-pass filter. B. Effect of contralateral level on CAP for different probe levels and subjects. The effect of a contralateral elicitor is plotted as a fractional change in dB (cf. insets Figs. 2C, 3C, 4C), with positive values indicating an increase in CAP amplitude in the presence *versus* absence of the elicitor. Different probe levels are shown with different symbols (inset). Subjects (n=5) are differentiated by color (the data of the subject in A are shown in dark brown). Single data points were obtained with paradigm SP1. Colored dashed lines connect data from same subject at the same elicitor level: these data were obtained with stimulus paradigm SP2. Black symbols and heavy line show averages across all datapoints; error bars indicate SEM. Probe frequency was 4 kHz.

Fig. 5B shows the influence of the sound level of the contralateral elicitor on the CAP response to a 4 kHz probe tone, for five subjects (different colors). The effect is shown as a fractional (dB) change in CAP amplitude; different symbols indicate different probe levels (Lprobe, see caption Fig. 5B). The response changes of the subject of Fig. 5A are shown in the dark brown color. Despite the intermittent presentation of the contralateral elicitor and the exploration of a wide range of elicitor and probe levels, effects of the contralateral elicitor remain small and inconsistent, both within and across subjects, and the spread in the data increases with elicitor level. Overall, most values trend positively, which is not the direction expected for this unmasked paradigm. The average across all data (Fig. 5B, thick black line with empty circles) does not significantly differ from 0 dB except for a small positive response change at an elicitor level of 70 dB (but note the lack of data points at 70 dB for subjects showing negative response changes: orange and purple).

### Electrophysiological effects of probe frequency

Recent evidence from human psychoacoustics and cochlear emissions points to an influence of the contralateral MOCR on low-frequency responses (see Discussion for references). This introduces an experimental challenge as it is well-known that CAPs are poorly suited to study low-frequency neural responses (Antoli-Candela and Kiang, 1978). Fortunately, at low frequencies another neural property (besides the onset synchronization required for CAPs) can be exploited to gain insight in neural responses, namely their ongoing synchronization to the fine-structure of the stimulus waveform. To explore the effects of a contralateral elicitor on low-frequency responses, we therefore studied the synchronized ongoing neural activity of the nerve, called the neurophonic (Weinberger et al., 1969; Snyder and Schreiner, 1984). This component is prominently present in the mass potential measured near the cochlea, but is entangled with receptor potentials. Particularly in view of the classical opposite effects of MOC activation on receptor potentials (increase) and neural responses (decrease), and the fact that these AC responses can sum constructively or destructively, it is important to separate the contributions of these two generators. We previously developed two paradigms to separate neural from receptor generators (Verschooten and Joris, 2014; Verschooten et al., 2015, 2018). A first paradigm exploits the property of neural adaptation – which is not present at the receptor level. Although it is the more sensitive and robust paradigm, it makes use of an ipsilateral forward masker and provides only a transient window on neural responses, which makes it less suited in the context of the present study. A second, less sensitive but simpler paradigm extracts the 2nd harmonic over the entire duration of the probe response to low-frequency probe tones. Previous work (Snyder and Schreiner, 1984; Forgues et al., 2013; Verschooten et al., 2015) shows that at the probe frequencies used here (≤ 800 Hz) this component is dominated by neural phase-locking rather than the receptor potential. Neural contributions are detectable at probe frequencies > 800 Hz but become more difficult to interpret because neural phase-locking to fine-structure declines to levels comparable to or smaller than the receptor potentials.

In two subjects, we measured the 2^nd^ harmonic derived spectrally from the response to long tones (800 ms) using stimulus paradigm SP3 (Fig. 1). Fig. 6 shows measurements for one subject. Fig. 6A illustrates envelope traces of the sharp bandpass-extracted 2nd harmonic obtained from responses to probe tones of 300 – 700 Hz at 80 dB in the absence (blue) or presence (red) of a contralateral elicitor also at 80 dB. At all these probe frequencies, a sustained reduction in 2^nd^ harmonic is present. The bottom trace shows responses to a 3-kHz probe tone, which are small and unaffected by the presence of the elicitor.

**Figure 6.**
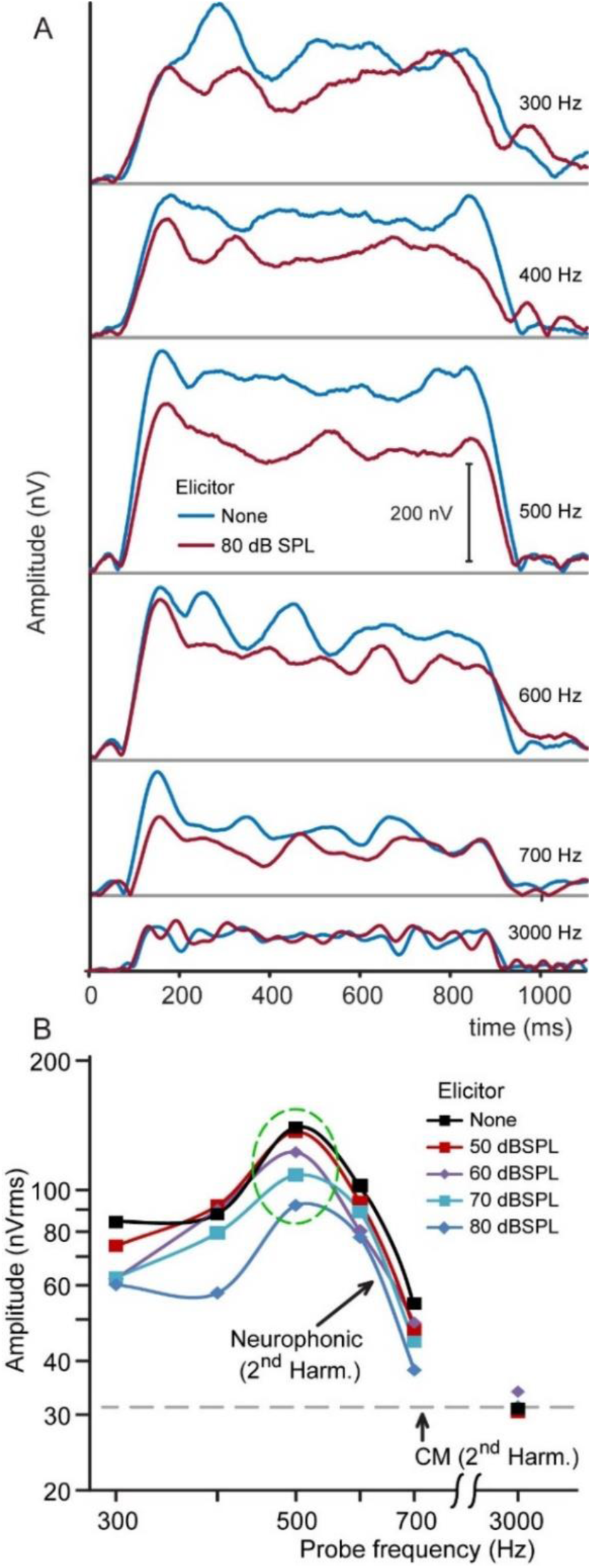
Low-frequency cochlear neural suppression. Examples of the influence of a contralateral elicitor on the magnitude of the neurophonic’s 2^nd^ harmonic, in one subject. The elicitor was a continuous Gaussian broadband (100Hz - 16kHz) noise at 80 dB SPL. A. Time course of the magnitude of the 2^nd^ harmonic in the neurophonic potential in response to 70-dB SPL probes at different frequencies (300 – 3000 Hz), in the presence (red) or absence (blue) of a contralateral elicitor. B. Magnitude of the 2^nd^ harmonic for different levels of the contralateral elicitor, including no-elicitor condition (black). Bottom curve (dark blue) is from series shown in A (80 dB SPL).

Fig. 6B shows the extracted 2nd harmonic amplitude of this subject for a range of probe frequencies and elicitor levels (none, 50, 60, 70 and 80 dB SPL). The magnitude contour of the neurophonic without contralateral elicitor (black) is consistent with that in previous studies (Verschooten et al., 2018, 2015; Verschooten and Joris, 2014): at low frequencies, the neurophonic increases with increasing frequency, then peaks (here at ∼500 Hz), to then strongly decline (700 Hz). Introduction of a contralateral elicitor (colored symbols) causes a reduction in amplitude which is progressive with elicitor sound level, for probe frequencies up to 700 Hz. This is particularly clear at 500 Hz (green dashed circle). At 3 kHz, the fine-structure in the response is entirely dominated by the receptor potential and there is no effect of the elicitor, independent of its sound level.

Data are replotted as a ratio (dB) as a function of level of the contralateral elicitor in Fig. 7 for two subjects. The dominant finding, illustrated with the trend averaged across probe frequencies (thick black curve with open circles), is an increasing reduction in probe response with increasing level of the contralateral elicitor. For an elicitor level of 80 dB, the average response changes were -3.2 dB (Fig. 7A) and -5.5 dB (7B). Linear fits of the average trendlines had slopes of respectively -0.08 dB/dB SPL (Fig. 7A) and -0.14 dB/dB SPL (Fig. 7B) and gave estimates of the contralateral elicitor threshold level (at 0 dB) of ∼ 45 dB SPL (Fig. 7A) and ∼ 35 dB SPL (Fig. 7B).

**Figure 7.**
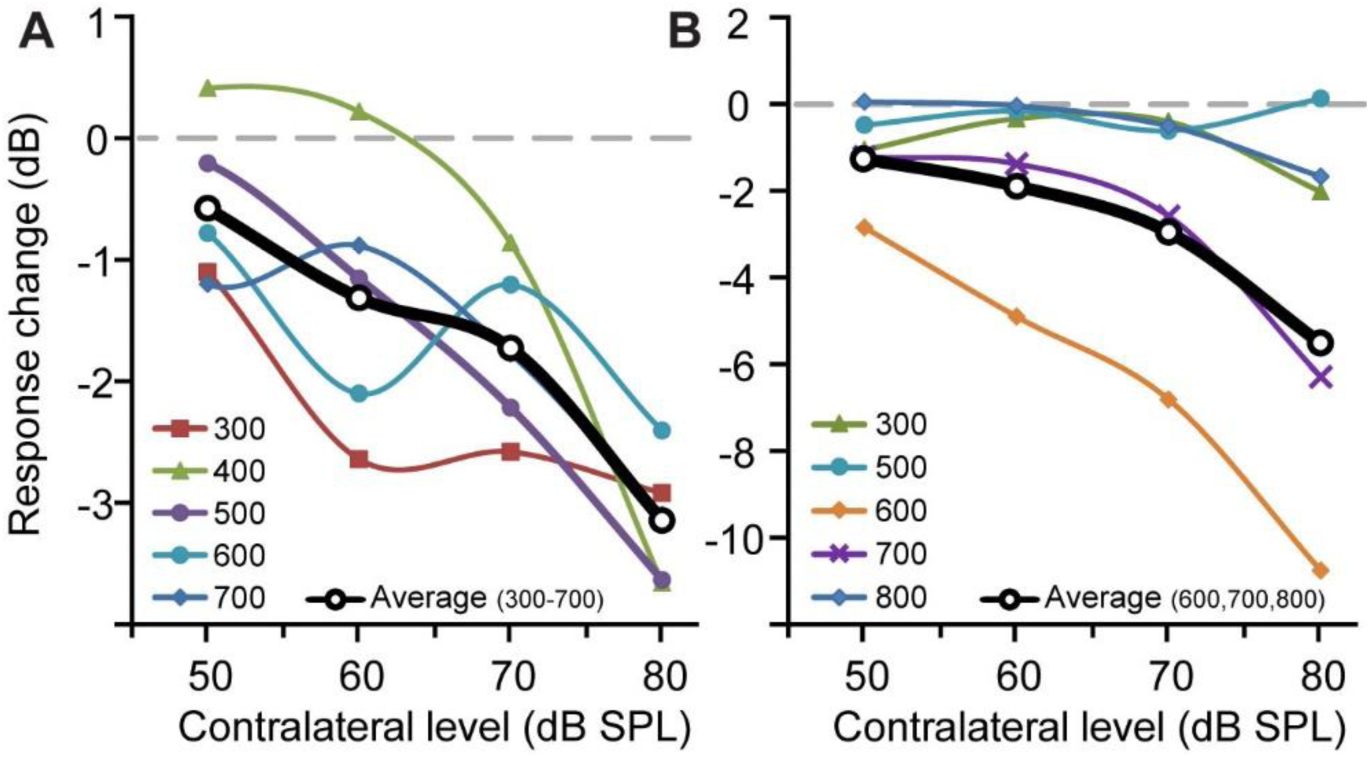
Increasing low-frequency cochlear neural suppression with increasing contralateral elicitor level, in two subjects. The change in (root-mean-square) amplitude of the 2^nd^ harmonic is shown as a function of elicitor level for different probe frequencies. The probe was a tone at 70 dB SPL. Response change is shown as the ratio (in dB) of the response in the presence of an elicitor relative to response (dashed line at 0 dB) in the absence of that elicitor. A,B: Data for two subjects. Curves through the data points are trendlines. Black curves represent the average of data across the specified frequencies.

In sum, whereas the study of CAP responses (with or without forward masking) revealed that a contralateral elicitor has only small and variable effects on probe responses of 4 and 6 kHz, the study of sustained neural responses to probe frequencies ≤ 800 Hz reveals more substantial and systematic suppressive effects. Note that the physiological data do not allow firm conclusions regarding the frequency range between 800 Hz and 4 kHz, due to the technical limitations of the method used. As stated above, the decrease in amplitude of neural *versus* receptor potentials limits the frequency range over which neural phase-locking can be examined *via* the second harmonic (more so than *via* the method of the “decaying neurophonic”: Verschooten et al. [2015]). The physiological results clearly suggest a low-frequency bias in suppressive neural effects of the contralateral MOCR. In the final section we examine whether behavioral measurements support this finding and whether we can better define the contralateral MOCR’s frequency range of operation.

### Psychophysical effects of elicitor level

We followed up with a simple psychophysical experiment in a group of 12 subjects. The expectation is that triggering the MOCR with an elicitor at one (the “contralateral”) ear increases the threshold in quiet for a probe tone at the other (“ipsilateral”) ear. We first investigated the influence of a contralateral elicitor on the detection threshold for a 4 kHz and 800 Hz tone, using stimulus paradigm SP4 (Fig. 1). Fig. 8A shows the change in threshold for five subjects tested with 4-kHz tones for different contralateral elicitor levels. All five subjects (different colors) showed a net increase in threshold over the range of elicitor levels studied. The average change in threshold (black) follows a monotonic increase with elicitor level, with a slope of 0.08 dB/dB SPL and a maximum threshold change of 3 dB at 80 dB SPL. The elicitor threshold level estimated by linear extrapolation is ∼45 dB SPL.

**Figure 8.**
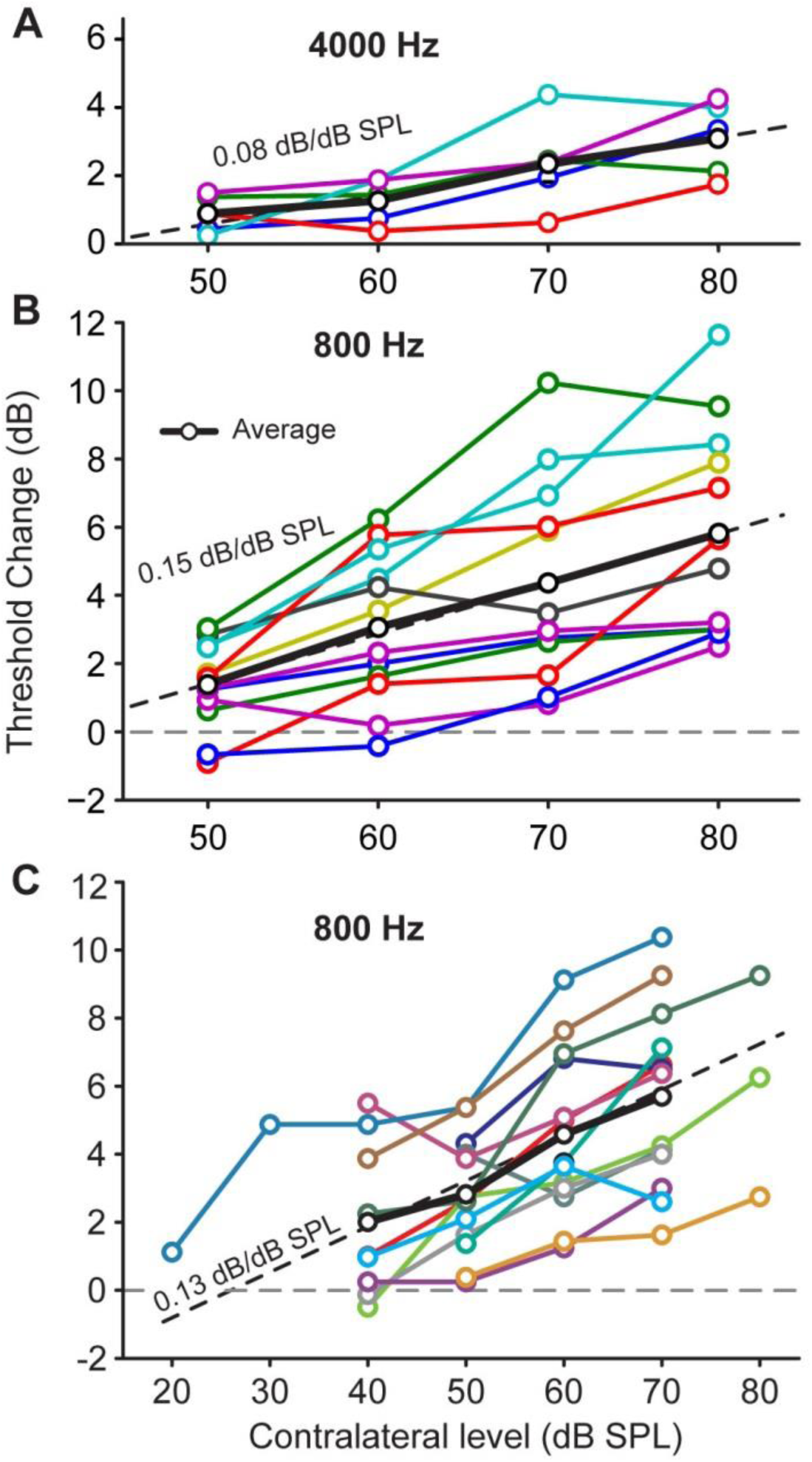
Elevation of detection threshold by a contralateral elicitor of increasing level. A: Effect of a contralateral noise elicitor on the threshold of hearing for an ipsilateral tone-pip of 4 kHz for different contralateral levels and subjects. The black line is the average of 5 subjects (colored lines); the slope is indicated by the dotted line. B: Same but for a tone-pip of 800 Hz and a larger group of subjects. C: Same as B but for a different group of subjects, using lower contralateral elicitor levels. In all panels, different colored lines represent different subjects. B and C are similar experiments but for different test groups. Only subjects that showed a threshold change were included.

Fig. 8B shows results for an 800-Hz probe tone in an extended subject group (n = 12, including 5 subjects tested at 4 kHz). The trend (black line) for the 800-Hz data is in the same direction as that at 4 kHz, but with ∼80% greater effect and with an average slope that is twice as large (0.15 dB/dB SPL *versus* 0.08 dB/dB SPL). The estimated threshold level of the elicitor, derived by linear extrapolation, is again ∼45 dB SPL.

The measurements at 800 Hz were repeated in a new group of 13 new subjects recruited to better define the frequency range of contralateral MOCR effects (see next section) where the range of elicitor levels used was lower (usually 40-70 dB). The results for this group are shown in Fig. 8C. The strength of the effect for different subjects is variable, but with similar slopes. The average slope of the trend (0.13 dB/dB SPL) for this group is very similar to that of the other group (Fig. 8B), but with a lower estimated threshold of ∼25 dB SPL. Across the two groups, we find some remarkable values, including a threshold change of up to 12 dB and a contralateral threshold level as low as 30 dB SPL.

The contralateral noise elicitor used for the experiments summarized in Fig. 8 was broadband and identical for all assessments. Early work in humans showed that the frequency content of the elicitor affects the strength of the contralateral MOCR (Folsom and Owsley, 1987) but this topic has been somewhat controversial (Lilaonitkul and Guinan, 2009, 2012; Aedo et al., 2015). In a few subjects we examined the influence of the elicitor spectrum (bandwidth and center-frequency) on the behavioral threshold to an 800 Hz tone, with the spectral level set at 60 dB SPL. Results for 3 subjects are shown in Fig. 8-suppl1, which depicts the elicitor bandwidth (abscissa) and the resulting change in threshold (ordinate). While in all cases broadband noise is the best MOCR elicitor, the threshold differences with noises that are band-limited to the frequency region of the probe are only a few dB or less.

### Psychophysical effects of probe frequency

Having established the large difference of contralateral effects at 4 kHz *versus* 800 Hz, we examined the probe frequency dependence of the MOCR in more detail. Fig. 9A-J shows threshold changes in 10 subjects for which relatively complete measurements were available. Despite intersubject variability in the size of the threshold increases in the presence of the contralateral elicitor, in all subjects the effect is larger at low frequencies and is undetectable at the highest probe frequency tested (6.4 kHz). Fig. 9K shows a summary scatterplot with all data except those of 2 subjects (subjects of Fig. 9I,J, who basically lacked an effect, even at 800 Hz). The trendline (red line; MATLAB RLOESS, range = 0.55) reveals a low-pass behavior with a cutoff at ∼4 kHz indicated by the vertical dashed line (intersection of extrapolated trendline with abscissa).

**Figure 9.**
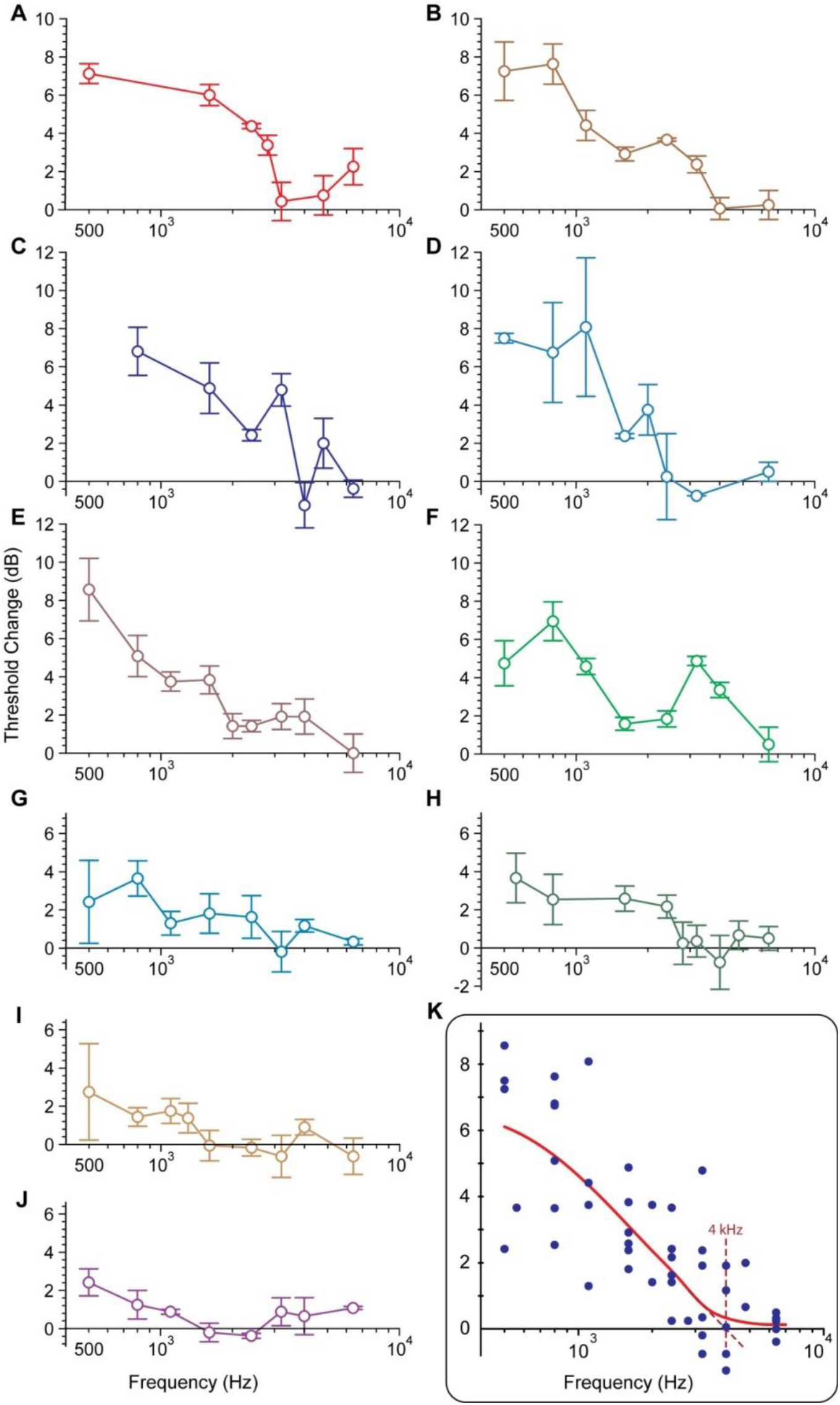
Threshold elevation by contralateral noise as a function of ipsilateral probe frequency, for ten subjects. A-J: Individual data. The contralateral level was 60 dB SPL, except for the results in D where it was 55 dB SPL due to an exceptional low MEMR threshold of 68 dB SPL. Each data point is the average threshold change with and without elicitor. Error bars represent propagated SEM. K: Population trend. Data from 8 subjects with a threshold increase of at least 2 dB at 800 Hz were included, excluding subjects of panels I,J.

### Comparison of psychophysical and electrophysiological results

The psychoacoustic experiments measured a change in behavioral threshold, while the physiological experiments measured an amplitude change in neural activity. Fig. 10 compares both sets of data for low probe frequencies (≤ 800 Hz) and 4 kHz probe tones by converting the behavioral threshold change into a “response change” (increase in threshold corresponds to a negative response change). The behavioral data in Fig. 10A,B (magenta lines) correspond to the trendlines of Figs. 8B and 8A, respectively, which were obtained for the same range of contralateral elicitor levels. The physiological data of Fig. 10A,B (green lines) are based on the averages of the data in Fig. 7A,B and Fig. 5B, respectively. A direct quantitative comparison between these behavioral and physiological results is of course fraught with difficulties: the comparison in Fig. 10 is simply meant as a shorthand to pull together the various datasets and to point out parallels in the behavioral and physiological observations. At low frequencies (Fig. 10A), a clear suppressive effect of the contralateral elicitor is evident both behaviorally and physiologically, which increases with elicitor level. At 4 kHz (Fig. 10B), no net suppressive effect was observed in physiological experiments (green); a suppressive change (magenta) was observed in the behavioral data but was not as marked as at low frequencies (red, Fig. 10A).

**Figure 10.**
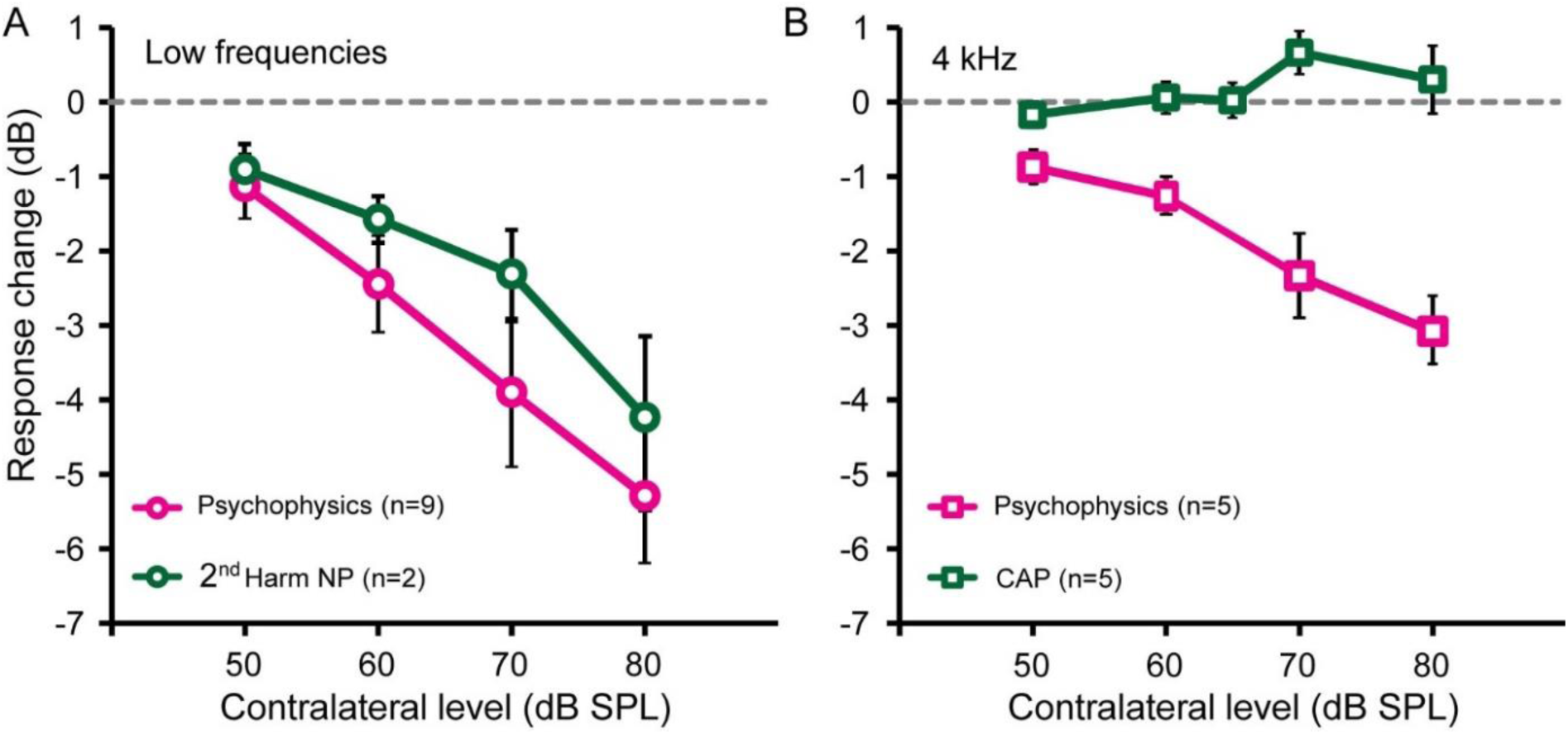
Comparison between psychophysical and physiological population trends obtained in this study. A: Effect of increasing contralateral elicitor level at low probe frequency (800 Hz). Data from Figs. 8B (magenta, behavior) and 7A,B (green, physiology, averaged across two subjects). B: Same, for a probe frequency of 4 KHz. Data from Figs. 8A (magenta, behavior) and 5B (green, physiology). For comparison, psychophysical results (threshold changes) are represented as response changes (negative dB values). Error bars represent SEM.

## Discussion

### Summary

We assessed the contralateral sound-evoked MOCR in awake, normal-hearing humans using psychophysical and physiological techniques. We first recorded mass-potentials in the middle ear using a transtympanic method and examined anti-masking effects of a sustained, broadband, contralateral elicitor on forward-masked CAP responses to brief tones at 4 and 6 kHz (Figs. 2-4). Effects were small and variable and were inconsistent across and within subjects. We modified and simplified the paradigm and conducted a parametric study in which we examined effects of a gated contralateral elicitor on CAP responses to 4-kHz probe tones, while parametrically varying the level of the probe and elicitor (Fig. 5). Again, effects observed were small and inconsistent across subjects. We then investigated the effect of the MOCR at much lower probe frequencies. This necessitated a different stimulus and analysis paradigm: measurement of the 2^nd^ harmonic of the sustained neurophonic response to a low-frequency probe tone enabled us to examine neural phase-locking to fine-structure. We found that a contralateral elicitor caused a clear reduction in neural response (Figs. 6,7), which was evident for probe frequencies up to 800 Hz and was absent at 3 kHz. For probe frequencies between these limits (800-3000 Hz) the 2^nd^ harmonic is difficult to interpret because of the decline in neural phase-locking relative to the receptor potentials (CM). We concluded with psychophysical measurements, partly on the same subjects who participated in the physiological study, which showed that detection thresholds were weakly affected by a contralateral broadband elicitor at 4-kHz tones (Fig. 8A), but more strongly at 800 Hz (Fig. 8B,C). Examination of this frequency dependence in further detail showed a population trend with a clear bias toward low frequencies for the contralateral MOCR, with an upper limit close to 4 kHz (Fig. 9K).

### Comparison to previous studies of contralateral stimulation on neural potentials in human

CAP suppression by contralateral sound is a rapid way to assess efferent effects in animals, and was suggested as a technique to investigate MOCR effects in humans (Liberman, 1989). Such efforts have given variable results. Most studies looked for contralateral suppression of mass potentials to clicks recorded at the tympanic membrane or ear canal. Large effects of tones at the contralateral ear were claimed in an early, short report (Folsom and Owsley, 1987) where the N1 potential was recorded in the ear canal in response to filtered (4 kHz) clicks, but concerns have been raised regarding its methodology (Puria et al., 1996; Lichtenhan et al., 2016). Much smaller and variable suppression effects, on average near 2 dB effective attenuation, were reported in 5 subjects by averaging potentials recorded in the ear-canal to clicks, over many (∼10) hours (Lichtenhan et al., 2016). Somewhat larger effective attenuation was found in response to clicks (up to 3 dB) and particularly in response to frequency chirps (up to 5 dB) (Smith et al., 2017). Najem et al. (2016) found that tonal contralateral stimuli suppressed CAPs at the tympanic membrane more for ipsilateral 1 and 4 kHz tone pips than for clicks but the suppressive effects were small (maximally 1/3 reduction in amplitude). Anti-masking effects of contralateral noise on CAPs in response to clicks for a group of 11 subjects were also small (1/3 to 1/4 of the fractional changes reported in cat)(Kawase and Takasaka, 1995). Large suppressive effects of contralateral noise were reported from 3 anesthetized patients who underwent decompression surgery for a neurovascular conflict (Chabert et al., 2002): the CAP recorded directly from the auditory nerve in response to clicks showed an effective attenuation of 10 dB.

### Low-frequency bias of contralateral MOCR

Our main conclusion is that efferent effects of contralateral stimulation are most convincingly observed at frequencies below a few kHz. By examining the neural contribution to phase-locked mass potentials, we could examine sustained effects in response to tones of long duration and low frequency (rather than examining the synchronized onset response to the transient, high-frequency part of short probe stimuli). Our finding of a low-frequency bias was not anticipated by the studies of human neural potentials cited in the preceding section, which could not address frequency-dependence of efferent effects due to various constraints in the methods used (click or chirp probes; use of CAP; low SNR necessitating long recording times). Studies of experimental animals and of human anatomy did not anticipate our finding either, but the low-frequency bias is in line with some human studies, particularly of OAEs.

A number of studies of OAEs reported stronger suppressive effects of contralateral stimulation at low than at high probe frequencies (Kim et al., 2002; Lilaonitkul and Guinan, 2009, 2012; Francis and Guinan, 2010; Xing and Gong, 2017), but a low-frequency bias is less clear for behavioral measures (Kawase et al., 2003; Marrufo-Pérez et al., 2021; Salloom and Strickland, 2021). The innervation density of MOC efferents, as estimated from labeling of their terminals, is higher in the basal half of the human cochlea, as it is in other animals (Liberman and Liberman, 2019). Adjusted for the frequency-position mapping across species (Greenwood, 1990), maximum innervation density in young human subjects was predicted to be at cochlear sites tuned near ∼2-4 kHz. The mismatch between this prediction and our finding of a low-frequency bias is surprising but there are many other circuit features that may be involved besides terminal density. It is unknown whether the cochlear distribution or projection patterns in human differ for the circuits underlying the contra- and ipsilateral MOCR, as is the case in animals (Guinan et al., 1984; Liberman, 1988; Brown, 2014). Also, it is clear from physiological studies in anesthetized animals (Liberman, 1988) that many factors can affect the strength of the contralateral MOCR for different probe frequencies (CF, threshold, suprathreshold response strength, binaurality, aftereffects), but how these results extrapolate to awake humans is unknown.

### Lack of physiological anti-masking effects at 4-6 kHz

Animals provide physiological evidence for the anti-masking effect of the MOCR: CAP responses to masked probe tones can increase in amplitude in the presence of a contralateral elicitor. In a detailed study in cat using contralateral noise elicitors (Kawase and Liberman, 1993), it was found that suppression of CAP responses to unmasked tones was largest for probe tones at low sound levels and low (2-8 kHz) frequencies, while the unmasking effect (in µV) was largest for probe tones with high sound levels and at high (8-16 kHz) frequencies. In some animals, contralateral stimulation caused enhancement of CAP responses to unmasked tones rather than suppression, particularly at high probe frequencies. This was argued to be an unmasking effect due to activation of the most sensitive nerve fibers by internally generated noise (e.g. from breathing). Possibly the small increases in CAP responses to unmasked and masked tones for the subject illustrated in Fig. 2A,C,E reflect such enhancement.

The variability in physiological effects at 4-6 kHz does not necessarily imply that there are no efferent effects in these subjects, only that our measurements could not provide convincing evidence for them. A case can be made that we observed antimasking of the CAP at 4 kHz in one subject (Fig. 2) and an elevation of detection threshold at that frequency in some of the subjects tested (e.g. Fig. 8A, 9F). Discrepancies between effects on single fibers and populations responses have been noted by others (Guinan, 2018). Efferent effects on single afferent neurons are not necessarily reflected in mass potentials, which only report on those behaviors that affect synchronized electrical responses. In cat, the suppressive effect of contralateral tones is strongest on auditory nerve fibers of low and medium spontaneous rate tuned near 1-2 kHz (Warren and Liberman, 1989b), which are not the main generators of the CAP (Versnel et al., 1990; Bourien et al., 2014). Nevertheless, for the stimulus conditions explored in our study, sizable unmasking effects are present for CAP measured in response to tone pips in anesthetized cat (Liberman, 1989; Kawase and Liberman, 1993) and would be expected to be stronger in unanesthetized subjects (Boyev et al., 2002; Guitton et al., 2004; Chambers et al., 2012; Aedo et al., 2015). The parallels between the perceptual and physiological trends (Fig. 10) also suggest that the lack of consistent efferent effects in the physiological measurements at higher probe frequencies (4 – 6 kHz) is not simply due to a limitation of the recording technique.

Overall, an unsatisfactory imbalance remains between the size of the purported efferent effects in humans and the methodological intricacies that seem to be needed to pull out these effects. Two recent salient observations are that cochlear efferent innervation in humans is sparse relative to experimental animals (Liberman and Liberman, 2019), and that MOC efferents are driven more by descending (Schofield and Beebe, 2019; Romero and Trussell, 2022) inputs than by ascending input from the cochlear nucleus (Romero and Trussell, 2021). Perhaps the anatomically more robust system in experimental animals enables its study in a state where it is only engaged by its peripheral input, while the sparser system in humans requires active engagement in an auditory task to reveal clear anti-masking effects.

## Materials and Methods

This study was carried out in accordance with the recommendations of good clinical practice (ICH/GCP) and in accordance with the Declaration of Helsinki, and approved by the Medical Ethics Committee of the University of Leuven (study S56783, protocol ECochG-EF-P-2). Written informed consent was obtained from all subject before the study commenced.

### Subjects

For the transtympanic recordings, volunteers were recruited via an advertisement on campus. The age of the subjects was between 20 and 30 years. Subjects were requested to avoid exposure to loud sounds in the days preceding the experimental session. On the morning of the experimental session or the day before, the subject’s hearing was assessed including an inquiry for hearing problems, an otological exam by an otolaryngologist, a pure tone audiogram (thresholds < 20 dB nHL, 125 Hz - 8 kHz), bilateral tympanometry to assess middle ear function, and measurement of the acoustically evoked threshold of the MEMR. The latter was determined at both sides for a 1 kHz tone and broadband noise (MADSEN, ZODIAC 901). A total of 6 subjects (3 male, 3 female) were tested for contralateral MOCR effects using transtympanic recording, of which 2 (1 male, 1 female) also participated in the psychophysical experiments. Additional subjects were recruited for psychophysical experiments: selection was identical to that for the physiological experiments except that no otological exam was performed. Here, 25 subjects passed the screening (6 male, 19 female). Participants in either study had the right to end their participation at any time.

The duration of the electrophysiological experiments varied between 1 and 4 hours. Experiments were conducted in a double-walled, soundproofed, and electrically shielded booth with a windowed door (Industrial Acoustics Company, Niederkrüchten, Germany). Subjects chose a comfortable reclined position on a bed and were asked to remain still during the recordings. Both the subject and experimenters were grounded to the booth’s potential using antistatic wrist straps to prevent electrical static discharge. During the actual experiment, an observer was present inside the booth to monitor the status of the subject and maintained visual contact with the experimenters outside the booth.

The psychophysical experiments lasted 2 to 6 hours per subject, depending on individual performance and progress. They were conducted in a double-walled, soundproof, and electrically shielded booth, similar to the one used for electrophysiological experiments. The booth was equipped with a comfortable chair, two insert earphones for acoustical stimulation, and computer equipment, including a screen, keyboard, and mouse connected to an external computer. A monitor outside the booth allowed the researcher to guide and oversee the experiment.

### Trans-tympanic procedure

Evoked mass potentials were recorded near the cochlea using an approach *via* the human middle ear using a trans-tympanic procedure, extensively described elsewhere (Verschooten and Joris, 2022). For every subject, a custom silicone earmold (Dentsply, Aquasil Ultra XLV regular) was created of the right ear. The mold contained two cast openings, each 2 mm wide, for calibration, visualization, needle insertion and acoustic stimulation. The entire acoustic system was calibrated *in situ* with a probe-microphone (Etymotic Research, ER-7C) positioned near the tympanic membrane. The earphone-speaker was connected to one of the openings of the earmold via a plastic T-piece, which also served as access port for a rigid endoscope with camera (R. WOLF, 8654.402 25 degree PANOVIEW; ILO electronic GmbH, XE50-eco X-TFT-USB) to visualize the ear canal and tympanic membrane. During acoustic calibration, the openings were sealed with acrylic impression compound (Audalin, Microsonic), except for a tiny opening in one tube to prevent static pressure build-up. Prior to electrode insertion, the tympanic membrane and ear canal were anesthetized locally with Bonain’s solution (a mix of equal amounts of cocaine hydrochloride, phenol and menthol), which was aspirated after 30 minutes. A short sterile plastic tube was inserted into the earmold to accommodate the sterile needle-electrode (TECA, monopolar disposable, 75mm x 26G, 902-DMG75-TP). Ground and reference electrodes were connected to the recording equipment. The needle-electrode was inserted and gently placed through the tympanic membrane and positioned on the cochlear promontory or in the round window niche under endoscopic guidance. To ensure stable positioning and good electrical contact, the needle-electrode was maintained under slight tension with rubber bands supported by a custom frame, which was positioned over the external ear and secured around the head with Velcro straps. Subjects typically reported a short-lasting and vague sensation of touch during electrode insertion. Once the electrode was in place, the tube openings were resealed with Audalin, and the electrode was connected to a preamplifier.

Recording sessions were terminated after 4 hours or earlier when the subject requested to stop. At the end of the experiment, the needle electrode and earmold were removed, and an otoscopic examination was performed. Subjects were requested to keep the ear dry for 10 days following the recording session. An otolaryngologist was available in the weeks after the experiment to address any concerns or conduct a follow-up examination.

### Psychophysical procedure

Subjects underwent a short training session to familiarize themselves with the experimental environment and paradigm. As the psychoacoustical tasks were demanding, measurements were divided into blocks, each separated by 15-minute breaks to help maintain concentration. Psychophysical detection thresholds, with and without contralateral sound, were obtained using a 3-alternative forced-choice (3AFC) method with a 2-down 1-up tracking procedure, implemented via a custom MATLAB (Mathworks, release 2014a) script (PsyAcoustX, Bidelman et al., 2015). The subject indicated the deviant interval using a mouse or keyboard. After completing a trial, the average detection threshold and standard deviation of the last eight inversions were calculated. To ensure data reliability, trials with a standard deviation exceeding 5 dB SPL were excluded. To address variability and potential time-dependent effects, measurements with the same conditions were repeated multiple times during the session.

### Acoustical stimulation

For the electrophysiological experiments, stimuli were generated using custom software and a digital sound system (Tucker-Davis Technologies, system 2, sample rate: 125 kHz/channel) consisting of a digital-to-analog converter (PD1), digitally controlled analog attenuators (PA5), a headphone driver (HB7) and electromagnetically shielded earphone-speakers (Etymotic Research, ER-2, 20 Hz - 16 kHz). The ipsilateral (re. side of recording, i.e. right ear) earphone-speaker was connected to the earmold with plastic tubing, while the contralateral earphone-speaker was connected to a disposable foam eartip (ER 1-14A, regular 13mm) inserted in the left ear canal. Ipsilateral stimuli were calibrated *in situ* with a probe-microphone (Etymotic Research, ER-7C).

For the psychoacoustical experiments, two ER-2 insert earphones with eartips were used for bilateral stimulation. These actuators have a nominally flat frequency response at the eardrum, as well as wide bandwidth, and high interaural attenuation. The earphones were driven by headphone amplifiers (TDT, HB7), connected to a LynxTWO sound card (Lynx Studio Technology, Inc.) installed in a computer outside the soundproof booth. The computer managed stimulus generation, acoustic stimulation, visualization, and experiment control. At the start of the session the acoustic system was calibrated at 1 kHz.

### Electrophysiological recording

Evoked potentials were measured using a low-noise differential preamplifier (Stanford Research Systems, SR560). On the (ipsilateral) side of the recording, the signal input was connected to the needle-electrode, the reference input to an earlobe clamp with conductive gel, and the ground input was connected to a standard disposable surface electrode at the mastoid. The battery-operated preamplifier was galvanically isolated (A-M systems, Analog stimulus isolator Model 2200) from the mains-powered equipment outside the sound booth. The signal was subsequently amplified (DAGAN, BVC-700A), band pass filtered (30 Hz - 30 kHz, 12 dB/octave cut-off slopes), and recorded (TDT, RX8, ∼100 kHz/channel, max. SNR 96 dB). Recorded data were stored and further analyzed using MATLAB. Stimuli and recorded signals were monitored in real time via an oscilloscope (LeCroy, WaveSurfer 24Xs).

### Stimulus paradigm

The stimulus design accommodates several, sometimes conflicting, requirements. It should enable assessment of effects on pure tones and masked pure tones; disambiguation of effects on receptor and nerve potentials; comparison of our results to previous electrophysiological and psychophysical work; and at the same time accommodate the limited time and quality typical for human recordings. Five paradigms (Fig. 1) were standardly used across subjects; additional parameters were explored when time allowed. The common goal of the different stimulus paradigms was to assess whether a contralateral broadband Gaussian white noise (the *elicitor*) affects the mass potential or the perception of an ipsilateral tone (the *probe*). Paradigms SP0 through SP3 were used in the physiological experiments, SP4 only in the psychophysical experiments. The stimulus levels never exceeded 95 dB SPL and were chosen to be below the subject’s threshold of the acoustic reflex (MEMR), as detailed below (section *Evaluation of MEMR threshold*).

In all physiological stimulus paradigms (SP0 – SP3), the ipsilateral probe alternated in polarity (Fig. 1: red and blue) to aid disentanglement of receptor and neural contributions. Such pairs of stimuli were presented with or without a contralateral elicitor to trigger the contralateral MOCR, and the main question of interest is whether response components are affected by the presence of the contralateral elicitor.

SP0 was a simple paradigm used to determine an appropriate probe level for further testing with SP1. Within a block, a 100-ms probe tone at the ipsilateral ear, fixed in frequency and alternating in polarity, is varied in SPL across a number of conditions (e.g. Fig. 2A: absence, 50, 60, 70, 80, 85 dB). In some blocks, a continuous elicitor was presented at the contralateral ear, while in others, it was absent. To minimize spectral splatter, all signals were gated with a 5 ms raised cosine. The probes were 100 ms long, including on- and off-ramping, and separated by 284 ms.

SP1 assessed anti-masking, i.e. whether MOCR-activation weakens the influence of a forward masker relatively more than it attenuates the probe. This is similar to a condition used by Kawase and Liberman (1993). It is the only paradigm that contains a masker (magenta), which is a Gaussian white noise. The basic stimulus unit consists again of pairs of probe tones (blue, red) that are identical except in polarity. They are preceded by a masker with an inter-stimulus pause of 5 ms (not including on- and off-ramping) to allow the offset-response to the masker to settle before the onset-response to the probe. The stimulus parameter of interest is masker SPL, which varied over a number of conditions (e.g. Fig. 2E: absence, 40, 50, 60, 70 dB). Within each experimental block, the masker is rotated through conditions (Fig. 1: masker 1, 2, …) for multiple repetitions (n=100). Blocks were alternated with and without a continuous contralateral elicitor (green), which was a continuous Gaussian broadband noise similar to the masker but with fixed level and bandwidth (100 Hz - 16 kHz). The whole procedure was then repeated for different probe and elicitor levels. A visual representation of block results is shown in Results (Figs. 2C, D).

SP2 and SP3 have no masker and have an intermittent elicitor. They differ in the length of the ipsilateral probe stimulus: SP2 uses a 100-ms tone to obtain the CAP and CM; SP3 uses a longer tone of 800 ms to enable measurement of the 2^nd^ harmonic component in the sustained response. Probe tones were fixed in frequency and level but alternated in polarity. The contralateral elicitor is activated 200 ms before the start of the probe and remains active until 20 ms after its end. The intervals between probes were ∼400 ms. Within a block, the contralateral elicitor varied in SPL across a number of conditions (e.g. Fig. 5B: absence, 50, 60, 70, 80 dB), indicated in Fig. 1 as noise 1, noise 2, etc. The temporal parameters were chosen, based on animal and human psychophysical experiments, to allow the MOC system to be activated by the onset of the elicitor and to recover between different conditions.

SP4 was used for behavioral testing with the 3AFC method where a pure-tone probe is randomly presented in one of three intervals. The method is described in Salloom and Strickland (2021). The task of the listener is to indicate the interval that contains the probe. Depending on the outcome, the intensity of the probe is increased (after one incorrect answer) or decreased (after two correct answers), which tracks 70.7% correct (Levitt, 1971). After 12 level reversals the trial automatically stops and the average threshold and standard deviation are calculated from the last eight reversals. The start intensity of the probe was typically 30 dB SPL. The duration of the probe was 12.5 ms for frequencies above 800 Hz and 25 ms otherwise, with 5-ms on and off gating. The elicitor was a 400-ms broadband white Gaussian noise (100-8000 Hz) which started 300 ms before the probe and had 20-ms on and off gating. One concern was a masking effect of the contralateral elicitor *via* interaural crosstalk. Control measurements with an ipsilateral masker, combined with the interaural attenuation characteristics of the Etymotic earphone-eartip combination, indicated that contralateral elicitor levels of 90 dB SPL or more were required to produce masking via crosstalk. Contralateral elicitors were restricted to levels ≤ 80 dB SPL.

### Evaluation of MEMR threshold

Ipsi- and contralateral stimulus levels were set below the subject’s threshold of the MEMR, determined during initial screening. Electrophysiological recording provided an additional check for MEMR activation, which was easily detected by the presence of muscle action potentials in real time on an oscilloscope and auditory monitor, and as well as in the offline analysis. Fig. 1-suppl1 shows an example of prominent muscle artifacts (red marks) in the CM-filtered response (black trace; more details in legend) during and after stimulation by the probe (blue trace, representing the CM envelope). An initial test was conducted at the beginning of each recording session to determine the MEMR threshold, constrained to a maximum of 95 dB SPL. All reported physiological data were verified offline to confirm the absence of muscle artifacts. Furthermore, the lack of a significant decrease in CM amplitude at high contralateral sound levels served as further confirmation of the absence of MEMR effects.

### Mass potential analysis

From the mass potentials, one onset and two sustained response components were extracted: the CAP, the CM and the 2nd harmonic of the ongoing response. The latter is a periodic signal and distortion product which has been shown in different species to be dominated by neural generators at low frequencies and by receptor potentials (CM) at high frequencies (e.g. Fig. 10 in Verschooten et al., 2015). In previous studies, we have shown a large neural contribution to the 1^st^ harmonic of the mass potential (Verschooten and Joris, 2014; Verschooten et al., 2015, 2018). The 1^st^ harmonic is multiple times larger than the 2nd harmonic, but disentanglement of the 1st harmonic from that of the CM requires a more complex paradigm based on adaptation (forward masking), which may unintentionally activate the ipsilateral MOCR (Verschooten et al., 2017). In contrast, the 2^nd^ harmonic is simpler to extract and can be measured over extended time spans, which favors its signal-to-noise ratio. We therefore limited ourselves here to the measurement of the 2^nd^ harmonic as an index of neural activity. Its amplitude is extracted from the same signal that contains the CAP, but unfiltered.

The CAP and 2^nd^ harmonic amplitude values were extracted from the averaged summed responses to stimulus conditions with opposite probe polarity; the CM was extracted from their difference (divided by 2). The SNR of the human responses is significantly smaller than in (smaller) laboratory animals. Reduction of the uncorrelated background noise is therefore required by averaging over many repetitions (n > 100). For the CAP, the summed responses were further filtered off-line with a non-causal low-pass filter based on the RLOESS function (MATLAB). The RLOESS input parameter “range” was chosen such that the spectral cutoff frequency was ∼3 kHz. The magnitude of the CAP was obtained between the first positive and first negative peak (P1-N1) (for example waveforms, see Fig. 5A). For the CM and the 2^nd^ harmonic, the amplitudes were calculated from the spectra, obtained with a fast Fourier transform (FFT, MATLAB). To minimize any on/offset phenomena, the parts of the probe responses 15 ms after onset and 5 ms before offset were excluded. The amplitudes were calculated as the root-mean-square (RMS) sum of the significant spectral components (> 3 times background noise) around the frequency of interest, i.e. the probe frequency for the CM and twice the probe frequency for the 2^nd^ harmonic.

### Statistical analysis

The central tendency (mean) of the CAP and CM in each measurement block was calculated Bootstrapping (n=400) was performed for each block to estimate variability, and the Standard Error of the Mean (SEM, shown as error bars in the Figures) was calculated. Additionally, a Shapiro-Wilk test confirmed normality for each block. For paradigms with repeated measurement blocks (i.e., SP0 and SP1), the mean was calculated across all repeated blocks for each condition. The SEM for these conditions was derived as the propagated SEM from these repeated blocks, rather than through bootstrapping all measurements. This approach mitigates the influence of time-related drift in central tendency on variance. Finally, a two-sample t-test with pooled variance was applied under the null hypothesis (μ1 = μ2), using a two-sided interval. Assumptions for the t-test, such as equal variance, were confirmed to ensure validity.

## Acknowledgments

The authors express their heartfelt gratitude to Dr. Jana Van Canneyt and Msc. Linde Peeters for their invaluable assistance, which significantly contributed to the success of this work. This research was supported by grants from BOF (OT-14-118 to PXJ) and NIH (R01 grant DC008327 to EAS).

## Author Contributions

*Conceptualization*: EV,EAS,PXJ; *Methodology*: EV; *Investigation*: EV,NV; *Software*: EV; *Validation*: EV,EAS,PXJ; *Formal analysis*: EV; *Resources*: EAS,PXJ; *Data curation*: EV; *Writing - original draft*: EV,PXJ; *Writing – Review & Editing*: PXJ,EV,EAS,NV; *Visualization*: EV; *Funding acquisition*: EAS,PXJ.

