## Supplementary material for "Assessment of Contralateral Efferent Effects in Human *Via* ECochG": Figure 1 Supplement 1

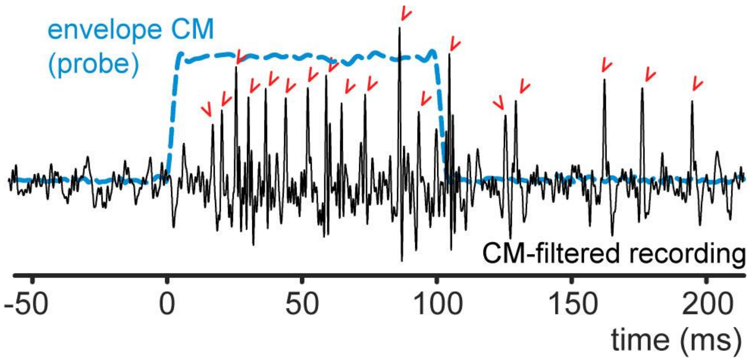


**Figure 1 Supplement 1**. Example of stapedius muscle artifacts in the recording during the activation of the acoustic reflex. The black and blue trace are derived from the same raw stimulus-evoked response to a 6 kHz probe tone at 85 dB SPL, but with different filtering. The blue dashed trace is the envelope of the cochlear microphonic (CM: Hilbert transform of band-pass filtered signal around the probe frequency of 6 kHz), which indicates the timing of the probe stimulus. The black trace is a low-pass filtered version of the raw response. Muscle artifacts are indicated by red check marks. The timeline (bottom) is synchronized to the onset of the probe tone. Note that the first artifact occurred ~15 ms after the onset of the CM.
