## Supplementary material for "Assessment of Contralateral Efferent Effects in Human *Via* ECochG": Figure 8 Supplement 1

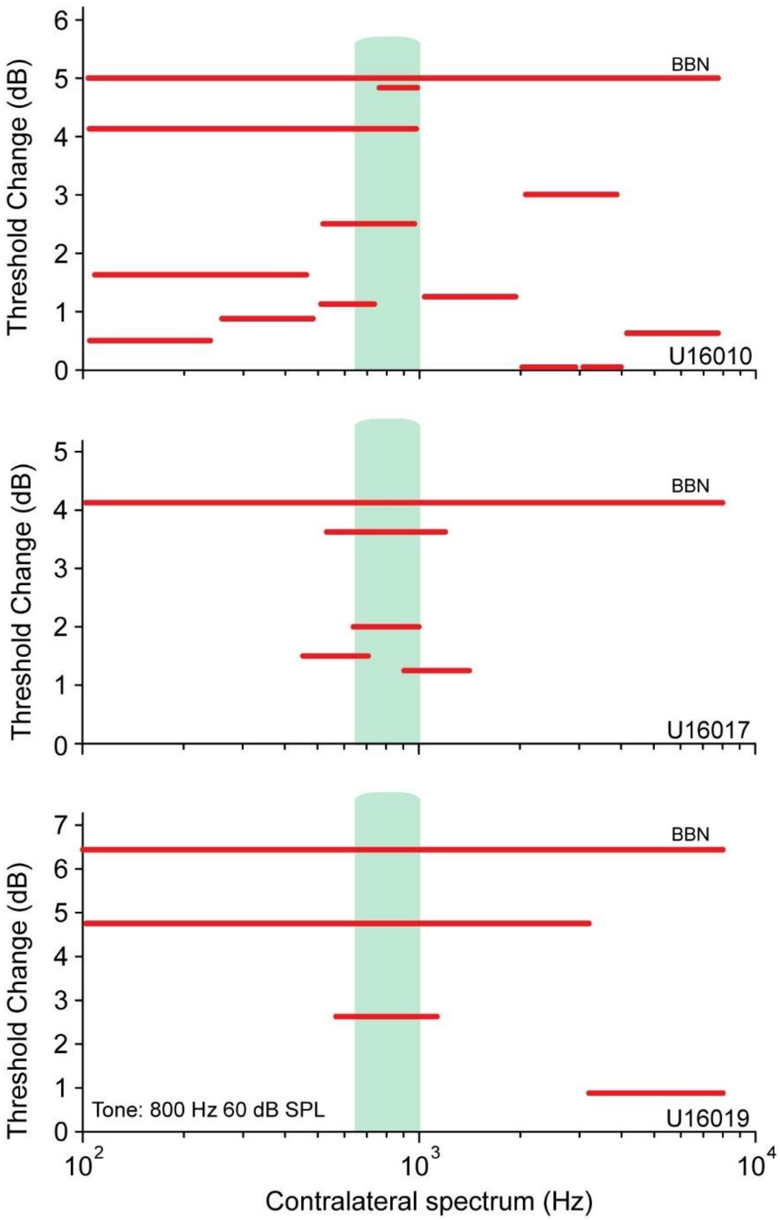


**Figure 8 Supplement 1.** Elicitor bandwidth effects in three subjects. Threshold changes for detection of an 800-Hz probe in the presence of elicitors with different noise spectra (red lines). The vertical position of the red lines indicates the threshold change; their horizontal extent indicates the elicitor's spectrum. The contralateral spectral level was 60 dB SPL. Green shading indicates a 1/3 octave bandwidth centered at 800 Hz, for reference.
